# Structural modeling supports an interaction between the *Drosophila* Estrogen-Related Receptor and Sima/HIF1α

**DOI:** 10.64898/2026.07.22.739904

**Authors:** Yuan Feng, Jason M. Tennessen

## Abstract

The *Drosophila* Estrogen-Related Receptor (dERR) is an orphan nuclear receptor that regulates developmental metabolism, yet the protein cofactors that modulate its activity remain poorly defined. Here, we used the AI-based FlyPredictome platform to identify candidate dERR interaction partners and evaluate their predicted structural interfaces. Among the highest-confidence interactors is the transcription factor Sima, which represents the *Drosophila* ortholog of HIF1α – an interaction that was reciprocally identified in both dERR and Sima/HIF1α interaction datasets. Structural modeling predicted binding between the dERR ligand-binding domain and a conserved LXXLL motif in Sima/HIF1α. Notably, the predicted dERR– Sima/HIF1α interaction interface includes residues previously shown to be essential for *in vitro* binding, providing independent structural support for this biologically relevant protein–protein interaction.

## INTRODUCTION

Nuclear receptors (NRs) constitute a large superfamily of transcription factors that regulate gene expression in response to hormonal, metabolic, and environmental cues (Jin et al., 2025, Taubenheim et al., 2021). These highly conserved proteins act as molecular sensors that translate extracellular and intracellular signals into coordinated transcriptional responses, thereby playing central roles in development, energy metabolism, reproduction, and physiological homeostasis across metazoans (Jin et al., 2025, Doering et al., 2022, Vanderhaeghen et al., 2022, Li et al., 2025, Love et al., 1983, Wu et al., 2023, Tao et al., 2020). Classical nuclear receptors are typically activated by endogenous small lipophilic ligands, including steroid hormones, thyroid hormones, and retinoids, which bind to the ligand-binding domain (LBD) and induce conformational changes that modulate DNA binding and recruitment of transcriptional cofactors (Jin et al., 2025, Tao et al., 2020).

In contrast, orphan nuclear receptors represent a subclass of the NR superfamily for which endogenous ligands have not been definitively identified (Jin et al., 2025). Rather than relying on ligand binding for activation, orphan nuclear receptors are frequently regulated through alternative mechanisms, including constitutive activity, post-translational modifications, and, importantly, protein–protein interactions with cofactors (Jin et al., 2025, Berger and Moller, 2002, Mohan and Heyman, 2003). Functionally, orphan nuclear receptors are involved in diverse biological processes, including metabolic regulation, developmental patterning, circadian rhythm control, and cellular stress responses (Jin et al., 2025, Safe et al., 2025, Yemanyi et al., 2023). Notably, these functions are highly conserved from invertebrates to mammals, underscoring their fundamental biological importance and evolutionary conservation (Jin et al., 2025, Yemanyi et al., 2023, Safe et al., 2025, Segraves, 1994, King-Jones and Thummel, 2005).

The *Drosophila melanogaster* Estrogen-Related Receptor (dERR) is the sole ortholog of the mammalian Estrogen-Related Receptor (ERR) family of orphan nuclear receptors (Tennessen et al., 2011, Fasteen et al., 2025, Zike et al., 2025). dERR has emerged as a critical regulator of metabolic reprogramming during development, particularly through its control of glycolytic metabolism during larval growth (Tennessen et al., 2011). Given its central role in metabolic regulation, identifying protein cofactors that interact with dERR is essential for understanding how its transcriptional activity is regulated in a context-specific manner. Moreover, elucidating these regulatory mechanisms has broader implications for human health, as ERR family members are widley implicated in metabolic diseases (Huss et al., 2015, Misra et al., 2016, Choi et al., 2022) and cancer (Ranhotra, 2015, Pradhan et al., 2026, Gulwani et al., 2023), where dysregulated energy metabolism is a hallmark phenotype (Hanahan and Weinberg, 2011). Thus, defining dERR-associated protein networks can provide mechanistic insight into conserved pathways, linking transcriptional regulation to metabolic homeostasis.

Despite its importance, systematic identification of nuclear receptor cofactors remains challenging. Traditional approaches, such as yeast two-hybrid screening, co-immunoprecipitation, and pull-down assays, have been widely used to detect protein– protein interactions (Rao et al., 2014). However, these methods are inherently low-throughput, labor-intensive, and often limited in their ability to capture transient or context-dependent interactions. In addition, they typically provide limited structural information regarding the interaction interface, which is critical for understanding the mechanistic basis of transcriptional regulation. These limitations highlight the need for complementary approaches that can efficiently predict and prioritize candidate interactions while also offering structural insights.

Recent advances in artificial intelligence (AI)–driven protein structure prediction have provided powerful new tools for addressing these challenges. FlyPredictome is a recently developed computational platform that leverages approximately 1.5 million pairwise predictions generated using AlphaFold-Multimer to infer protein–protein interactions in *Drosophila* (Kim et al., 2026). By integrating structural modeling with quantitative confidence metrics, including the interface Local Interaction Score (iLIS), FlyPredictome enables proteome-wide prediction and prioritization of candidate interactions. This structure-informed, high-throughput approach is particularly well suited for identifying previously unrecognized cofactors of orphan nuclear receptors.

To identify potential cofactors of dERR, we queried the FlyPredictome database and ranked candidate interaction partners based on their iLIS values. Consistent with previous biochemical studies of the ERR family, dERR itself emerged as the highest-scoring interactor, reflecting a known ability of this nuclear receptor family to form homodimers (Vanacker et al., 1999, Greschik et al., 2002, Horard et al., 2004, Giguère, 2008). In addition, Sima, the sole *Drosophila* ortholog of HIF1α, was identified as a top-ranked candidate interactor. This prediction was particularly noteworthy because dERR and Sima are known to physically interact *in vitro* and genetic studies suggest that these two transcription factors cooperatively regulate metabolic gene expression. Since the structural basis of dERR-Sima/HIF1α binding has not been fully characterized, we focused our computational analysis on this interaction and used FlyPredictome to model the molecular interface between the two proteins. The resulting model both supports and expands upon previous experimental findings by identifying a putative interaction interface that encompasses residues known to be required for dERR-Sima/HIF1α binding.

## RESULTS AND DISCUSSION

To identify potential interacting partners of dERR, we queried the FlyPredictome database and ranked candidate proteins based on their iLIS values. Consistent with prior biochemical evidence, dERR itself emerged as the top-scoring interaction partner, reflecting its known ability to form homodimers (Vanacker et al., 1999, Greschik et al., 2002, Horard et al., 2004, Giguère, 2008; Table S1). This result provides internal validation of the predictive framework and supports the reliability of the platform in identifying biologically relevant interactions. Among the high-confidence candidates, Sima, the sole *Drosophila* ortholog of mammalian hypoxia-inducible factor 1α (HIF1α), was identified as a prominent interactor, with an iLIS score of 0.600, placing it within the top 1% of predicted interactions (Table S1). Notably, this prediction was reciprocal, as dERR also ranked among the top 1% of predicted interactors for Sima/HIF1α (Table S2). These results are consistent with a significant interaction between dERR and Sima/HIF1α.

Intriguingly, previous studies have demonstrated that Sima/HIF1α can physically interact with dERR (Li et al., 2013), and that *sima* mutant animals exhibit metabolic and developmental defects during embryonic and larval stages that closely resemble those observed in *dERR* mutants (Heidarian et al., 2025). The predicted dERR–Sima/HIF1α interaction is therefore consistent with a biologically relevant mechanism that may underlie the overlapping phenotypes observed in the two mutant backgrounds.

To further characterize the predicted dERR–Sima/HIF1α interaction, we examined the structural interface generated by FlyPredictome (Figure 1A–D). Interface mapping localized the interaction to the ligand-binding domain of dERR and a conserved LXXLL motif within Sima/HIF1α. Notably, LXXLL motifs are well-established nuclear receptor-binding motifs that interact with the activation function-2 (AF-2) surface of receptor ligand-binding domains (Savkur and Burris, 2004, Plevin et al., 2005). This binding arrangement is highly consistent with previous studies demonstrating a physical association between dERR and Sima/HIF1α (Li et al., 2013). Specifically, deletion of the C-terminal 12 amino acids of the dERR LBD (Figure 1C) abolishes Sima/HIF1α binding in vitro (Li et al., 2013), indicating that this region is required for complex formation. Likewise, substitution of the terminal leucine residues within the Sima/HIF1α LXXLL motif (Figure 1C) with alanines disrupts binding to dERR, demonstrating the functional importance of this motif (Li et al., 2013). Together, these findings show that the FlyPredictome model accurately recapitulates structural determinants previously demonstrated to mediate dERR– Sima/HIF1α binding, providing independent support for the biological relevance of the predicted complex.

**Figure 1.**
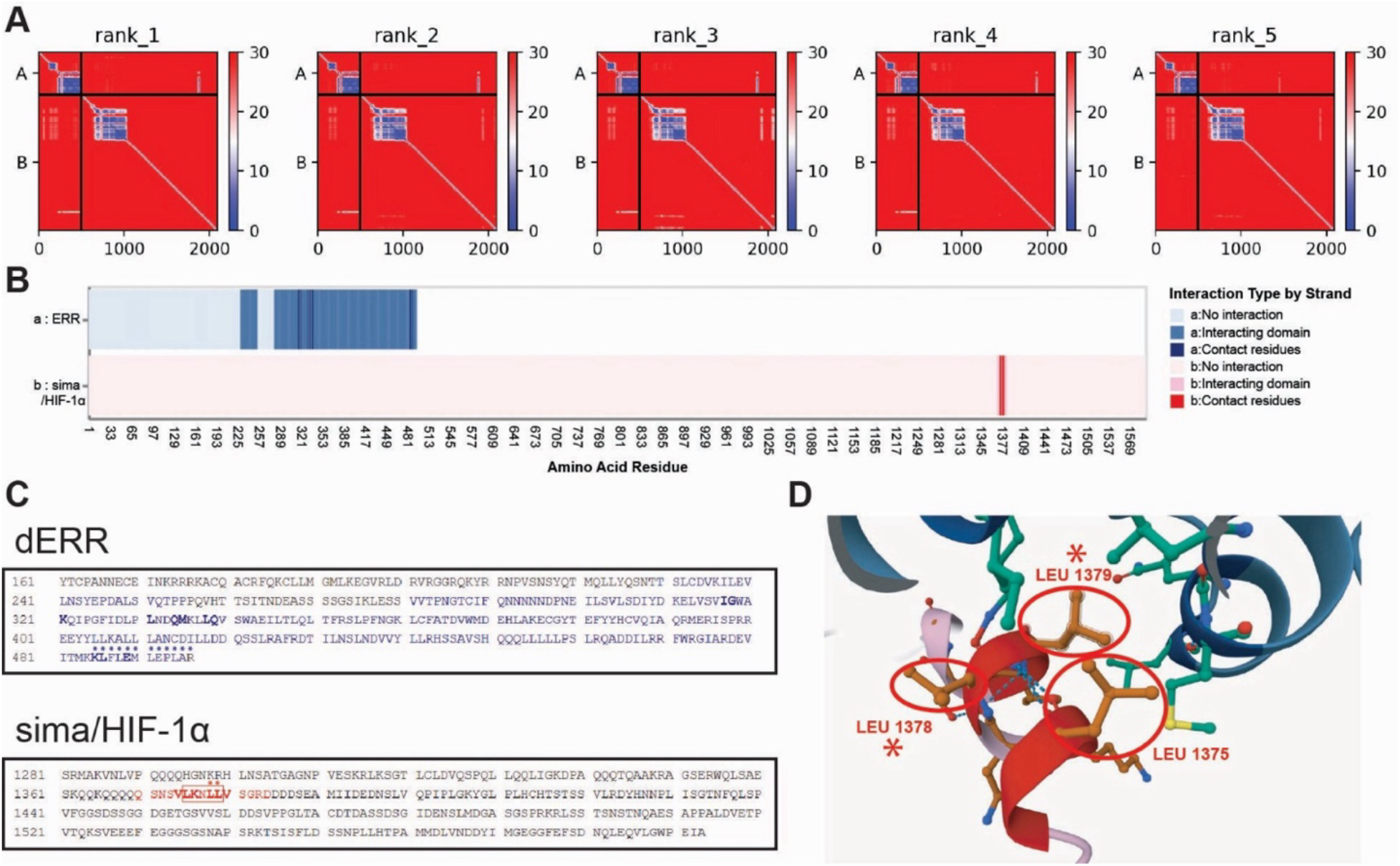
The *Drosophila* Estrogen-Related Receptor (dERR) is predicted to bind Sima/HIF1α by FlyPredictome. Predicted Aligned Error (PAE) heatmap of dERR and Sima/HIF1α interaction. (A) Each cell in the PAE heatmap represents the expected positional error between pairs of residues; low values (dark blue) indicate high confidence in their relative positioning. Off-diagonal blue regions between dERR and Sima/HIF1α indicate a confidently predicted interaction interface. Predictions are ranked from 1 to 4 according to confidence, with rank 1 representing the highest-confidence model. (B) Predicted interacting interface between dERR and Sima/HIF1α. The top bar represents the dERR protein sequence, and the bottom bar represents the Sima/HIF1α protein sequence. Light blue indicates the predicted interaction region in dERR, with dark blue highlighting contact residues. In Sima/HIF1α, the interaction region is shown in pink, with contact residues highlighted in red. (C) Predicted interacting amino acid sequence between dERR and Sima/HIF1α. Similar to (B), dark blue indicates the predicted interaction region in dERR, with contact amino acid residues highlighted in bold. Amino acid residues previously shown to influence dERR–Sima/HIF1α binding in yeast two-hybrid assays are marked by dark blue asterisks (Li et al., 2013). In Sima/HIF1α, the interaction region is shown in red, with contact residues similarly indicated in bold. The red box marks the LXXLL motif, and amino acid residues previously shown to influence dERR–Sima/HIF1α binding in yeast two-hybrid assays (Li et al., 2013) are marked by red asterisks. (D) Representative 3D structures of the interaction interface between dERR and Sima/HIF1α. Amino acid residues are colored according to the same scheme as in (B). The three leucine residues within the Sima/HIF1α LXXLL motif are highlighted with red circles and labeled according to their positions in the protein. The mutated sequences in the yeast two-hybrid experiment (Li et al., 2013) are marked by the red asterisks.

In this study, FlyPredictome identified Sima/HIF1α as a high-confidence candidate interactor of dERR and predicted an interaction interface that recapitulates residues previously shown experimentally to be required for binding. These findings demonstrate how AI-based structural prediction can both discover candidate protein interactions and provide mechanistic insight into their molecular basis, thereby facilitating the identification and prioritization of biologically relevant regulatory networks. Several limitations of the FlyPredictome platform should nevertheless be considered. First, the predictive models do not account for post-translational modifications, such as phosphorylation, acetylation, or ubiquitination, which can influence protein conformation, stability, and interaction dynamics (Li et al., 2026; Lee et al., 2023). Second, the computational framework does not incorporate cellular or environmental context, including factors such as pH, compartmentalization, or molecular crowding, all of which can affect protein behavior in vivo (O’Brien et al., 2012; Fuentes-Lemus and Davies, 2023; Shaffer et al., 2005). Third, the predictions are limited to binary interactions and therefore do not capture the higher-order complexes that often mediate nuclear receptor function (Millard et al., 2013; Jin et al., 2025; Tao et al., 2020). Despite these limitations, FlyPredictome successfully recovered a previously documented dERR–Sima/HIF1α interaction and identified an interface that encompasses residues known experimentally to be required for binding. These results highlight the utility of AI-based structural approaches for generating mechanistic hypotheses and prioritizing biologically meaningful regulatory interactions for future study.

## METHODS

The protein interactors of dERR and Sima/HIF1α were predicted using the FlyPredictome platform (Kim et al., 2026). The identifier for dERR (FBgn0035849) and Sima/HIF1α (FBgn0266411) were obtained from FlyBase (release FB2026_02) (Öztürk-Çolak et al., 2024) and used as input for pairwise interaction analysis. FlyPredictome employs AlphaFold2-based (Jumper et al., 2021) multimer modeling to generate structural predictions of protein complexes and provide confidence metrics. Default thresholds for positive interactions were applied (iLIS > 0.223; LIS > 0.168; cLIS > 0.298; ipTM > 0.48; confidence score > 0.494).

The predicted dERR–Sima/HIF1α complex exhibited an interface local interaction score (iLIS) of 0.600 and an interface predicted TM-score (ipTM) of 0.55, indicating a high-confidence interaction. Multiple ranked models were generated, and the top-ranked model (rank 1 based on ipTM score) was selected for downstream analysis. Structural visualization was performed using a Mol*-based viewer (Sehnal et al., 2021), and the predicted complex was exported as a PDB file for further analysis and figure preparation.

## ACKNOWLEDGEMENTS

We thank Flybase (NIH 5U41HG000739) for being an invaluable resource for this study. This study was supported by the NIGMS of the National Institute of Health under award R35GM119557 to J.M.T.

## SUPPLEMENTAL TABLE LEGENDS

**Table S1: FlyPredictome prediction of dERR interactors**. Default thresholds for positive interactions were applied (iLIS > 0.223; LIS > 0.168; cLIS > 0.298; ipTM > 0.48; confidence score > 0.494).

**Table S2: FlyPredictome prediction of Sima/HIF1α interactors**. Default thresholds for positive interactions were applied (iLIS > 0.223; LIS > 0.168; cLIS > 0.298; ipTM > 0.48; confidence score > 0.494).

## Notes

### Competing Interest Statement

The authors have declared no competing interest.

## LITERATURE CITED

Berger, J. & Moller, D. E. 2002. The mechanisms of action of PPARs. Annu Rev Med, 53, 409–35.

Choi, J., Oh, T. G., Jung, H. W., Park, K. Y., Shin, H., Jo, T., Kang, D. S., Chanda, D., Hong, S., Kim, J., Hwang, H., Ji, M., Jung, M., Shoji, T., Matsushima, A., Kim, P., Mun, J. Y., Paik, M. J., Cho, S. J., Lee, I. K., Whitcomb, D. C., Greer, P., Blobner, B., Goodarzi, M. O., Pandol, S. J., Rotter, J. I., Fan, W., Bapat, S. P., Zheng, Y., Liddle, C., Yu, R. T., Atkins, A. R., Downes, M., Yoshihara, E., Evans, R. M. & Suh, J. M. 2022. Estrogen-Related Receptor γ Maintains Pancreatic Acinar Cell Function and Identity by Regulating Cellular Metabolism. Gastroenterology, 163, 239–256.

Doering, K. R. S., Cheng, X., Milburn, L., Ratnappan, R., Ghazi, A., Miller, D. L. & Taubert, S. 2022. Nuclear hormone receptor NHR-49 acts in parallel with HIF-1 to promote hypoxia adaptation in Caenorhabditis elegans. Elife, 11.

Fasteen, T. D., Hernandez, M. R., Policastro, R. A., Sterrett, M. C., Zenter, G. E. & Tennessen, J. M. 2025. The Drosophila estrogen-related receptor promotes triglyceride storage within the larval fat body. J Lipid Res, 66, 100815.

Fuentes-Lemus, E. & Davies, M. J. 2023. E_ect of crowding, compartmentalization and nanodomains on protein modification and redox signaling - current state and future challenges. Free Radic Biol Med, 196, 81–92.

GiguèRe, V. 2008. Transcriptional control of energy homeostasis by the estrogen-related receptors. Endocr Rev, 29, 677–96.

Greschik, H., Wurtz, J. M., Sanglier, S., Bourguet, W., Van Dorsselaer, A., Moras, D. & Renaud, J. P. 2002. Structural and functional evidence for ligand-independent transcriptional activation by the estrogen-related receptor 3. Mol Cell, 9, 303–13.

Gulwani, D., Upadhyay, P., Goel, R., Sarangthem, V. & Debraj Singh, T. 2023.Unfolding of Imminent Bio-Signatures in the Prognosis of Thyroid Cancer; The Emergence of Estrogen Related Receptor Gamma (ERRγ) as a Hurricane. Asian Pac J Cancer Prev, 24, 375–387.

Hanahan, D. & Weinberg, R. A. 2011. Hallmarks of cancer: the next generation. Cell, 144, 646–74.

Heidarian, Y., Fasteen, T. D., Mungcal, L., Buddika, K., Mahmoudzadeh, N. H., Nemkov, T., D’Alessandro, A. & Tennessen, J. M. 2025. Hypoxia-inducible factor 1α is required to establish the larval glycolytic program in Drosophila melanogaster. Mol Metab, 93, 102106.

Horard, B., Castet, A., Bardet, P. L., Laudet, V., Cavailles, V. & Vanacker, J. M. 2004. Dimerization is required for transactivation by estrogen-receptor-related (ERR) orphan receptors: evidence from amphioxus ERR. J Mol Endocrinol, 33, 493–509.

Huss, J. M., Garbacz, W. G. & Xie, W. 2015. Constitutive activities of estrogen-related receptors: Transcriptional regulation of metabolism by the ERR pathways in health and disease. Biochim Biophys Acta, 1852, 1912–27.

Jin, P., Duan, X., Huang, Z., Dong, Y., Zhu, J., Guo, H., Tian, H., Zou, C. G. & Xie, K. 2025. Nuclear receptors in health and disease: signaling pathways, biological functions and pharmaceutical interventions. Signal Transduct Target Ther, 10, 228.

Jumper, J., Evans, R., Pritzel, A., Green, T., Figurnov, M., Ronneberger, O., Tunyasuvunakool, K., Bates, R., žíDek, A., Potapenko, A., Bridgland, A., Meyer, C., Kohl, S. A. A., Ballard, A. J., Cowie, A., Romera-Paredes, B., Nikolov, S., Jain, R., Adler, J., Back, T., Petersen, S., Reiman, D., Clancy, E., Zielinski, M., Steinegger, M., Pacholska, M., Berghammer, T., Bodenstein, S., Silver, D., Vinyals, O., Senior, A. W., Kavukcuoglu, K., Kohli, P. & Hassabis, D. 2021. Highly accurate protein structure prediction with AlphaFold. Nature, 596, 583–589.

Kim, A. R., Comjean, A., Veal, A., Rodiger, J., Han, M., Hu, Y. & Perrimon, N. 2026. FlyPredictome: A structural atlas of predicted protein-protein interactions in Drosophila. bioRxiv.

King-Jones, K. & Thummel, C. S. 2005. Nuclear receptors--a perspective from Drosophila. Nat Rev Genet, 6, 311–23.

Lee, J. M., HammaréN, H. M., Savitski, M. M. & Baek, S. H. 2023. Control of protein stability by post-translational modifications. Nat Commun, 14, 201.

Li, Q., Zhou, X., Zhang, X., Zhang, C. & Zhang, S. O. 2025. Nuclear receptor signaling regulates compartmentalized phosphatidylcholine remodeling to facilitate thermosensitive lipid droplet fusion. Nat Commun, 16, 3955.

Li, W., Wei, Q., Llanos, M., Gathmann, C., Governa, P., Chiu, T. Y., Wozniak, J. M., Jadhav, A. M., Holcomb, M., Cravatt, J., Dongre, A., Huang, M. L., Forli, S. & Parker, C. G. 2026. Posttranslational modifications remodel proteome-wide ligandability. Nat Chem Biol.

Li, Y., Padmanabha, D., Gentile, L. B., Dumur, C. I., Beckstead, R. B. & Baker, K. D. 2013. HIF- and non-HIF-regulated hypoxic responses require the estrogen-related receptor in Drosophila melanogaster. PLoS Genet, 9, e1003230.

Love, C. A., Cowan, S. K., Laing, L. M. & Leake, R. E. 1983. Stability of the human nuclear oestrogen receptor: influence of temperature and ionic strength. J Endocrinol, 99, 423–33.

Millard, C. J., Watson, P. J., Fairall, L. & Schwabe, J. W. 2013. An evolving understanding of nuclear receptor coregulator proteins. J Mol Endocrinol, 51, T23–36.

Misra, J., Kim, D. K., Jung, Y. S., Kim, H. B., Kim, Y. H., Yoo, E. K., Kim, B. G., Kim, S., Lee, I. K., Harris, R. A., Kim, J. S., Lee, C. H., Cho, J. W. & Choi, H. S. 2016. O-GlcNAcylation of Orphan Nuclear Receptor Estrogen-Related Receptor γ Promotes Hepatic Gluconeogenesis. Diabetes, 65, 2835–48.

Mohan, R. & Heyman, R. A. 2003. Orphan nuclear receptor modulators. Curr Top Med Chem, 3, 1637–47.

O’Brien, E. P., Brooks, B. R. & Thirumalai, D. 2012. E_ects of pH on proteins: predictions for ensemble and single-molecule pulling experiments. J Am Chem Soc, 134, 979–87.

ÖZtüRk-ÇOlak, A., Marygold, S. J., Antonazzo, G., Attrill, H., Goutte-Gattat, D., Jenkins, V. K., Matthews, B. B., Millburn, G., Dos Santos, G., Tabone, C. J. & Consortium, F. 2024. FlyBase: updates to the Drosophila genes and genomes database. Genetics, 227.

Plevin, M. J., Mills, M. M. & Ikura, M. 2005. The LxxLL motif: a multifunctional binding sequence in transcriptional regulation. Trends Biochem Sci, 30, 66–9.

Pradhan, J., Samal, A. P., Khatoon, U., Prusty, M. & Elangovan, S. 2026. Estrogen-related receptor α in breast cancer: From molecular insights to targeted therapy. Biochim Biophys Acta Rev Cancer, 1881, 189525.

Ranhotra, H. S. 2015. Estrogen-related receptor alpha and cancer: axis of evil. J Recept Signal Transduct Res, 35, 505–8.

Rao, V. S., Srinivas, K., Sujini, G. N. & Kumar, G. N. 2014. Protein-protein interaction detection: methods and analysis. Int J Proteomics, 2014, 147648.

Safe, S., Oany, A. R., Tsui, W. N., Lee, M., Srivastava, V., Upadhyay, S., Hailemariam, A., Farkas, E., Kakwan, S., Kearns, C. & Sivaram, G. 2025. Orphan nuclear receptor transcription factors as drug targets. Transcription, 16, 224–260.

Savkur, R. S. & Burris, T. P. 2004. The coactivator LXXLL nuclear receptor recognition motif. J Pept Res, 63, 207–12.

Segraves, W. A. 1994. Steroid receptors and orphan receptors in Drosophila development. Semin Cell Biol, 5, 105–13.

Sehnal, D., Bittrich, S., Deshpande, M., Svobodová, R., Berka, K., Bazgier, V., Velankar, S., Burley, S. K., KočA, J. & Rose, A. S. 2021. Mol* Viewer: modern web app for 3D visualization and analysis of large biomolecular structures. Nucleic Acids Res, 49, W431–w437.

Shaffer, K. L., Sharma, A., Snapp, E. L. & Hegde, R. S. 2005. Regulation of protein compartmentalization expands the diversity of protein function. Dev Cell, 9, 545–54.

Tao, L. J., Seo, D. E., Jackson, B., Ivanova, N. B. & Santori, F. R. 2020. Nuclear Hormone Receptors and Their Ligands: Metabolites in Control of Transcription. Cells, 9.

Taubenheim, J., Kortmann, C. & Fraune, S. 2021. Function and Evolution of Nuclear Receptors in Environmental-Dependent Postembryonic Development. Front Cell Dev Biol, 9, 653792.

Tennessen, J. M., Baker, K. D., Lam, G., Evans, J. & Thummel, C. S. 2011. The Drosophila estrogen-related receptor directs a metabolic switch that supports developmental growth. Cell Metab, 13, 139–48.

Vanacker, J. M., Bonnelye, E., Chopin-Delannoy, S., Delmarre, C., CavaillèS, V. & Laudet, V. 1999. Transcriptional activities of the orphan nuclear receptor ERR alpha (estrogen receptor-related receptor-alpha). Mol Endocrinol, 13, 764–73.

Vanderhaeghen, T., Timmermans, S., Watts, D., Paakinaho, V., Eggermont, M., Vandewalle, J., Wallaeys, C., Van Wyngene, L., Van Looveren, K., Nuyttens, L., Dewaele, S., Vanden Berghe, J., Lemeire, K., De Backer, J., Dirkx, L., Vanden Berghe, W., Caljon, G., GhesquièRe, B., De Bosscher, K., Wielockx, B., Palvimo, J. J., Beyaert, R. & Libert, C. 2022. Reprogramming of glucocorticoid receptor function by hypoxia. EMBO Rep, 23, e53083.

Wu, G., Baumeister, R. & Heimbucher, T. 2023. Molecular Mechanisms of Lipid-Based Metabolic Adaptation Strategies in Response to Cold. Cells, 12.

Yemanyi, F., Bora, K., Blomfield, A. K. & Chen, J. 2023. Retinoic Acid Receptor-Related Orphan Receptors (RORs) in Eye Development and Disease. Adv Exp Med Biol, 1415, 327–332.

Zike, A. B., Abel, M. G., Fleck, S. A., Dewitt, E. D. & Weaver, L. N. 2025. Estrogen-related receptor is required in adult Drosophila females for germline stem cell maintenance. Dev Biol, 524, 132–143.

